# A modular microscale granuloma model for immune-microenvironment signaling studies *in vitro*

**DOI:** 10.1101/2020.04.14.040048

**Authors:** Samuel B. Berry, Maia S. Gower, Xiaojing Su, Chetan Seshadri, Ashleigh B. Theberge

**Author notes:** correspondence: Dr. Ashleigh Theberge.

## Abstract

Tuberculosis (TB) is one of the most potent infectious diseases in the world, causing more deaths than any other single infectious agent. TB infection is caused by inhalation of *Mycobacterium tuberculosis* (Mtb) and subsequent phagocytosis and migration into the lung tissue by innate immune cells (e.g., alveolar macrophages, neutrophils, dendritic cells), resulting in the formation of a fused mass of immune cells known as the granuloma. Considered the pathological hallmark of TB, the granuloma is a complex microenvironment that is crucial for pathogen containment as well as pathogen survival. Disruption of the delicate granuloma microenvironment *via* numerous stimuli, such as variations in cytokine secretions, nutrient availability, and the makeup of immune cell population, can lead to an active infection. Herein, we present a novel *in vitro* model to examine the soluble factor signaling between a mycobacterial infection and its surrounding environment. Adapting a newly developed suspended microfluidic platform, known as Stacks, we established a modular microscale infection model containing human immune cells and a model mycobacterial strain that can easily integrate with different microenvironmental cues through simple spatial and temporal “stacking” of each module of the platform. We validate the establishment of suspended microscale (4 μL) infection cultures that secrete increased levels of proinflammatory factors IL-6, VEGF, and TNFα upon infection and form 3D aggregates (granuloma model) encapsulating the mycobacteria. As a proof of concept to demonstrate the capability of our platform to examine soluble factor signaling, we cocultured an *in vitro* angiogenesis model with the granuloma model and quantified morphology changes in endothelial structures as a result of culture conditions (P < 0.05 when comparing infected vs. uninfected coculture systems). We envision our modular *in vitro* granuloma model can be further expanded and adapted for studies focusing on the complex interplay between granulomatous structures and their surrounding microenvironment, as well as a complementary tool to augment *in vivo* signaling and mechanistic studies.

## Introduction

Tuberculosis (TB) is one of the most potent infectious diseases in the world, causing more deaths than any other single infectious agent.(World Health Organization 2019) TB infection is caused by inhalation of *Mycobacterium tuberculosis* (Mtb) and subsequent phagocytosis and migration into underlying lung tissue and the lymph system by responding immune cells (e.g., alveolar macrophages, dendritic cells).(Saunders and Britton 2007; Kaufmann 2004) Due to the inability of these innate immune cells to clear Mtb, persistent Mtb induce an adaptive immune response, leading to the formulation of a fused mass of immune cells around Mtb known as a granuloma.(Saunders and Britton 2007) Within the granuloma, Mtb typically enter a latent phase characterized by a non-proliferative phenotype and lipid uptake(Russell et al. 2009; Peyron et al. 2008; Deb et al. 2009), leading to a latent infection that is effectively contained.(World Health Organization 2019) However, disruption of the delicate equilibrium between proinflammatory factors (e.g., tumor necrosis factor-α (TNFα), interferon-γ (IFNγ)), microenvironment conditions (e.g., hypoxia, pH), and immune cell populations (e.g., macrophages, T cells) can lead to reactivation of latent Mtb and deterioration of the granuloma, initiating an active TB infection and dissemination of infectious mycobacteria.(Russell et al. 2009; Ramakrishnan 2012)

Deciphering the impact of microenvironment variations around the granuloma remains a significant challenge, and researchers often rely on *in vivo* animal models or biological samples (e.g., blood, tissue biopsy), considered the gold standard for studying TB, to reconstruct this complex environment. These methods have laid the foundation for understanding the pathogenesis and immunology behind TB, yet many existing *in vivo* models do not accurately recapitulate Mtb infection as seen in humans (although recent advances in mouse models and the established zebrafish/*M. marinum* model are closing this gap)(Cronan et al. 2018; Myllymäki, Bäuerlein, and Rämet 2016; Gern et al. 2017; Cronan and Tobin 2014; Zhan et al. 2017; Yong, Her, and Chen 2018). Additionally, despite recreating the complexity of an *in vivo* environment, spatial manipulation and probing of the granuloma microenvironment through introduction or removal of immune and tissue components is difficult in most animal models.(Scanga and Flynn 2014; Foreman et al. 2017; Zhan et al. 2017) Further, human-derived biological samples provide detailed cellular information regarding the granuloma, the immune response, and disease status (Darboe et al. 2019; Guyot-Revol et al. 2006; Ogongo et al. 2020; Berry et al. 2010), but are inherently limited as they only reflect a singular point in time, rather than the dynamic interactions that occur during the early stages of infection or disease progression.

Alternatively, researchers have utilized *in vitro* models to examine specific processes and immune phenomena associated with TB infection and granuloma formation, augmenting the valuable information elucidated from *in vivo* models.(Peyron et al. 2008; Deb et al. 2009; Birkness et al. 2007; Kapoor et al. 2013; Elkington et al. 2019) These models, which often rely on infection of patient-derived peripheral blood mononuclear cells (PBMCs) with mycobacterial strains, have successfully mimicked granuloma formation and behavior through soluble factor signaling between immune cells (Birkness et al. 2007; Puissegur et al. 2004; Crouser et al. 2017), Mtb reactivation(Kapoor et al. 2013), and PBMC differentiation.(Peyron et al. 2008) However, many of these *in vitro* models consist of granulomas grown inside of well plates(Peyron et al. 2008; Birkness et al. 2007; Kapoor et al. 2013; Puissegur et al. 2004; Crouser et al. 2017), limiting the ability of the researchers to easily manipulate the microenvironment of the granulomas and increase the complexity of their granuloma models through multiculture and introduction of key components of the microenvironment on demand. Recently, more complex, biomimetic models have been developed that have successfully recapitulated important biological phenomena(Venkata Ramanarao Parasa et al. 2014; Venkata R. Parasa et al. 2017) and examined novel therapeutic approaches to combat TB infection (Tezera et al. 2017; Bielecka et al. 2017), while simultaneously demonstrating innovative and tractable platforms. However, these models face limitations in studies where users wish to subject granulomas to various microenvironmental cues over time, or in enabling the addition of tissue components after the model is established.

Building upon the foundation created by previous *in vitro* models, we present the creation of a novel microscale *in vitro* granuloma model that can be adapted to study the soluble factor signaling between granulomas and their surrounding microenvironment immediately following infection. Using a recently developed modular microfluidic coculture platform, known as “Stacks”(Yu et al. 2019), we demonstrate a multi-layered coculture that can be spatially and temporally manipulated to mimic different microenvironments and timepoints. The Stacks platform utilizes suspended cultures, wherein a droplet is contained in a well consisting of walls but lacking a ceiling or floor (Casavant et al. 2013; Humayun, Chow, and Young 2018; Berthier et al. 2019), thereby enabling users to vertically stack layers containing different cell types and place them in signaling contact.(Yu et al. 2019) The modular component of the Stacks, as well as of other microfluidic platforms, offers a notable advantage as users can optimize model conditions individually and connect each component to create different complex systems.(Yu et al. 2019; Ong et al. 2019) As a proof of concept, we use a model mycobacterial strain known to induce granuloma formation *in* vitro(Puissegur et al. 2004; Seitzer and Gerdes 2003), *Mycobacterium bovis* Bacillus Calmette-Guerin (BCG), with human blood-derived immune cells and validate its ability to form an *in vitro* granuloma model on the microscale (4 μL culture volume) in a layer of the Stacks platform. Further, to demonstrate the ability of our stackable microscale infection model to signal with its surrounding microenvironment, we miniaturize an existing *in vitro* angiogenesis model (containing primary human endothelial cells) within a separate stackable layer.(Yu et al. 2019; Koh et al. 2008; Lonza 2018) Here, we validate the development of our microscale *in vitro* granuloma model and demonstrate the capability of the system to support soluble factor signaling between the granuloma model and a separate stackable endothelial culture. We envision our modular *in vitro* granuloma model can be further developed to include additional layers of immune cells, tissue models, and pathogens for studies examining the complex interplay between granulomatous structures and their surrounding environment, as well as a complementary tool to augment *in vivo* signaling and mechanistic studies.

## Results and Discussion

### Microscale Granuloma Model Design and Overview

We present a modular *in vitro* platform that we adapted to enable the ability to add, modify, and manipulate the granuloma microenvironment for studying the effects of cellular signaling on granuloma formation and development. To create this *in vitro* model, we adapted a previously described open microfluidic platform (“Stacks”(Yu et al. 2019)) (Figure 1) that relies on key fluidic principles, namely capillary pinning, to enable vertical stacking and removal of discrete cell culture wells without leakage or horizontal flow between stacked layers. The pinning of fluids within this platform is vital to contain cultures within the open wells and allows for the connection and separation of the wells without bonding or disruption of the cultures, respectively. Additionally, the Stacks platform provides numerous advantages such as pipette accessibility (due to its open culture wells), bio- and imaging compatibility (due to its fabrication from polystyrene or polypropylene), and microscale culture wells. Further, the Stacks device relies on surface tension and capillary forces for functionality, removing the need for external pumps commonly associated with microfluidic chips (i.e., syringe pumps) and allowing it to fit within common cell culture materials (e.g., OmniTray™, petri dish) and incubators. For our *in vitro* model, we created two independent layers that can be clicked together to initiate paracrine signaling or separated for independent analysis, thereby allowing us to temporally introduce different signaling microenvironments to our *in vitro* granuloma model (Figure 1). (Yu et al. 2019) The first layer, herein called the granuloma layer, consists of an infection model of monocyte-derived macrophages (MDMs) and a model mycobacterium strain, *Mycobacterium bovis* Bacillus Calmette-Guérin (BCG), suspended in a 3D extracellular matrix (ECM) plug to mimic some aspects of *in vivo* granuloma behavior (e.g., pathogen encapsulation, soluble factor secretion, aggregate formation) previously observed in other *in vitro* granuloma models.(Birkness et al. 2007; Kapoor et al. 2013; Crouser et al. 2017) The second layer, herein called the endothelial layer, consists of an *in vitro* angiogenesis model in which endothelial cells are cultured on a hydrogel plug. By placing the primary human endothelial cells on a separate Stacks layer, our model can be used to examine the induction of angiogenic processes around the granuloma layer and how those angiogenic processes are affected by the soluble factor signaling profile of the granuloma layer. It is important to note that in our platform, the granuloma layer is not vascularized directly, as is observed *in* vivo(Datta et al. 2015), and that in our system, our endothelial layer is more akin to modeling the surrounding vasculature that is manipulated during early TB infection (Figure 1). (Polena et al. 2016)

**Figure 1:**
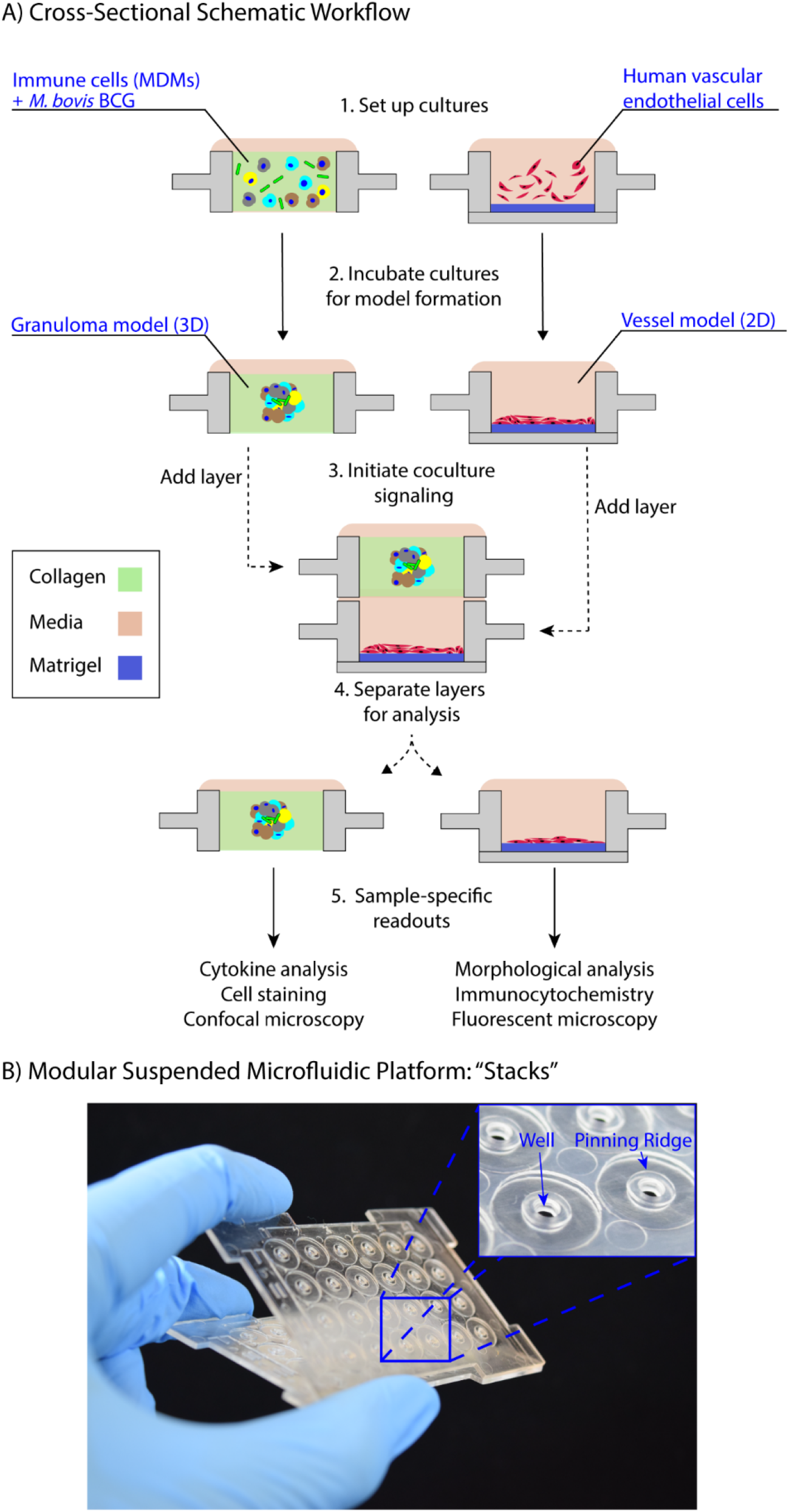
Suspended open microfluidic platform enables creation of a modular *in vitro* granuloma model. A) Schematic workflow showing cross sections of establishing the granuloma and endothelial layers and stacking of layers to initiate coculture paracrine signaling. B) The “Stacks” platform(Yu et al. 2019) contains an array of 24 individual suspended wells to facilitate the exchange of signals through the top and bottom of the well to neighboring layers and can be easily combined and removed. The inset illustrates the open suspended wells (2 mm diameter) and pinning ridges required to prevent leakage within the platform.

Previous *in vitro* granuloma models successfully recapitulated important components of granulomatous infections including leukocyte recruitment and signaling, establishment of dormancy and resuscitation, and genetic diversity at scales ranging from 12 well plates to 96 well plates.(Birkness et al. 2007; Crouser et al. 2017; Kapoor et al. 2013) Our model adds to these existing techniques through miniaturization and introduction of modularity to enable examination of the signaling phenomena between a mycobacterial infection and its surrounding microenvironment. We reduce the volume of our cultures (4 μL/well) more than 10-fold from previous examples (50-500 μL/well) to decrease cell and reagent usage in our model; further, we mix BCG and MDMs together in a 3D extracellular matrix (ECM), without pre-infecting the MDMs, to establish the infection in our granuloma layer after a minimum of 24 hours (Figure 1). However, due to the scale of the model (4 μL/well), the media buffering capacity, nutrient availability, waste accumulation, and multiplicity of infection (MOI) all needed to be optimized to permit successful formation of the granuloma layer (Supplementary Information 2). This is consistent with prior investigations on the effects of miniaturization on mammalian cultures.(Su et al. 2013) Similarly, we adapted and miniaturized the established *in vitro* angiogenesis assay(Lonza 2018; Koh et al. 2008) that is seeded into a separate Stacks layer and subsequently clicked together with the granuloma layer to initiate soluble factor signaling between the two layers (Figure 1, 4, Supplementary Information 3).

### Validation of Granuloma Model in Stacks Platform

To validate the successful establishment of a microscale *in vitro* granuloma model within our platform, we used three separate previously reported readouts: 1) aggregate formation, 2) encapsulation of the mycobacterium within host immune cells, and 3) soluble factor analysis.(Birkness et al. 2007; Kapoor et al. 2013; Crouser et al. 2017; Tezera et al. 2017) After initiating infection by mixing MDMs and BCG into the ECM (collagen I) and seeding it into the wells, we consistently observed aggregate formation in the granuloma layer containing BCG when wells were fixed and imaged on Day 4 post infection (p.i.) (Figure 2). Using mCherry-expressing BCG, we were able to observe aggregation of CellTracker Green-stained MDMs around the BCG in infection wells, whereas little to no aggregate formation was observed in the uninfected control wells (Figure 2); the 3D structure of the aggregates containing MDMs and BCG was confirmed through confocal imaging of the granuloma layers on Day 4 p.i. (Figure 2). We observed complete encapsulation of BCG within the multi-cellular aggregate, oftentimes noting the presence of multiple spatially distinct BCG within one aggregate and little to no extracellular BCG (Figure 2). An advantage of mixing the BCG and MDMs without direct preinfection of the MDMs is that MDMs must sample and migrate through the 3D collagen matrix in order to initiate the infection and respond to other infected MDMs, a process which is further supported by the microscale culture wells and an optimized MOI of 0.05.

**Figure 2:**
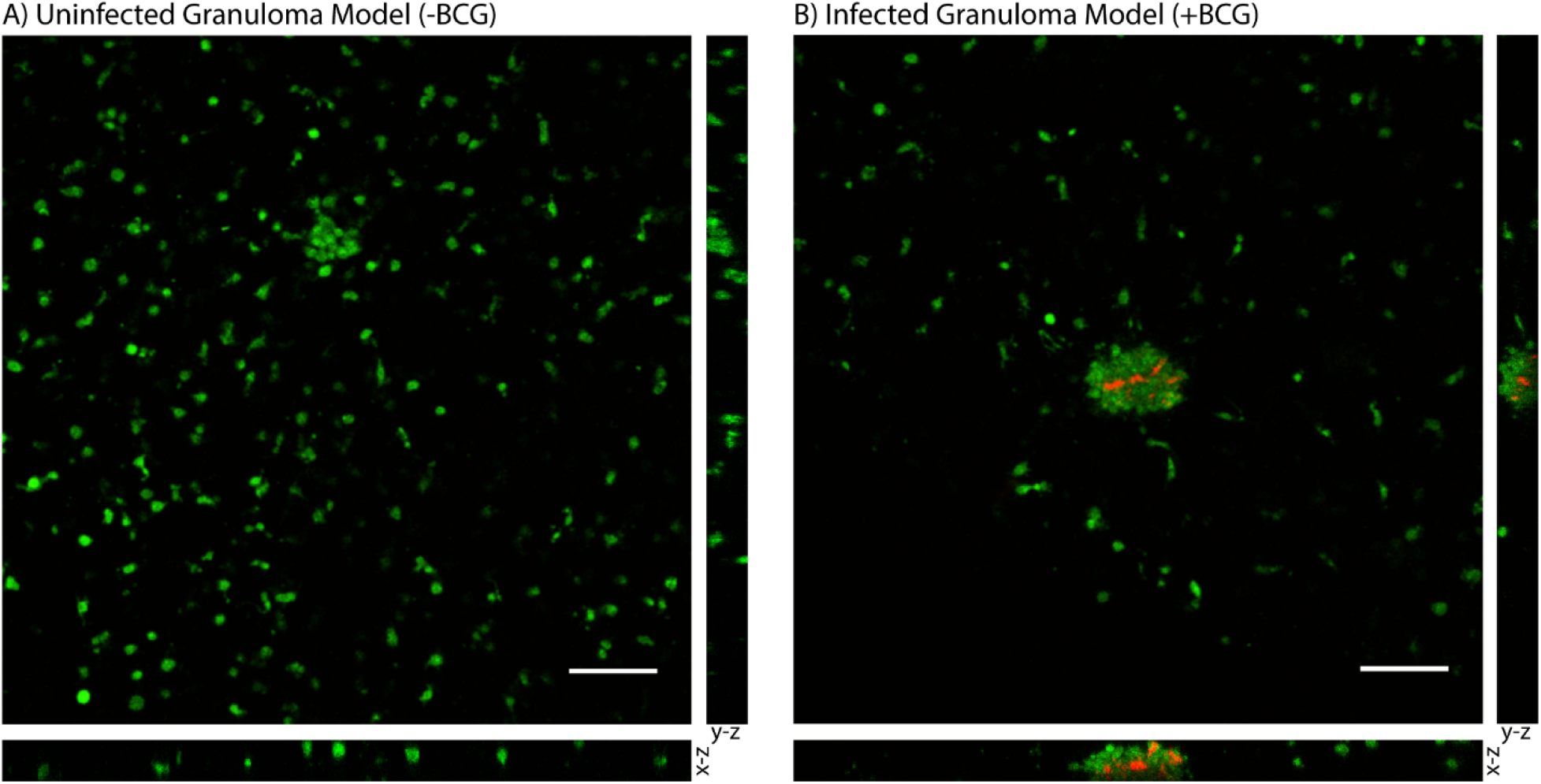
Aggregation of *M. bovis* BCG and MDMs in 3D collagen plugs within a layer of the Stacks microscale culture system. A) CellTracker Green-stained MDMs remain dispersed in a 3D collagen plug in the absence of coincubation with BCG. B) MDMs encapsulate mCherry-expressing BCG in the granuloma layer and form large multicellular aggregates that surround the BCG-infected MDMs and contain the mycobacterium within the aggregate. Confocal microscopy shows the structure of the aggregated MDMs with BCG and illustrates containment of the BCG within the aggregate of MDMs in the suspended 3D collagen plug. Representative images obtained on Day 4 p.i. MOI: 0.05. Scale bar: 100 μm. Bottom image depicts x-z plane (66.5 μm thickness) and right image depicts y-z plane (66.5 μm thickness).

To illustrate the capability of our model to be used for soluble factor signaling studies, we analyzed the secretion profile of three granuloma-associated proinflammatory factors: interleukin-6 (IL-6), tumor necrosis factor α(TNF α), and vascular endothelial growth factor (VEGF) (Figure 3).(Polena et al. 2016; Martinez, Mehra, and Kaushal 2013; Lin et al. 2007; Singh and Goyal 2013) In accordance with previous models and studies(Birkness et al. 2007; Polena et al. 2016; Martinez, Mehra, and Kaushal 2013; Singh and Goyal 2013; Lin et al. 2007) we observed significantly greater secretion of IL-6 (P = 0.005) and VEGF (P = 0.039) in our infected granuloma layers (+BCG) when compared to our control layers containing uninfected MDMs in monoculture (-BCG); further, we observed a 5-17-fold increase in TNFα secretion in 3 out of 4 independent experiments under the same conditions and a strong trend towards greater TNFα secretion (Figure 3). We also observed decreasing secretion of IL-6, TNFα, and VEGF over 5 days, with the greatest concentrations observed for IL-6 and TNFα on Day 1 and for VEGF on Day 2 (Figure 3). These results indicate that there is a large burst of proinflammatory factors immediately following infection, that then decreases and stabilizes over time as we begin to observe aggregate formation in the granuloma layers. Additionally, while we observe the anticipated increases in the secretion of these factors, we would expect a more robust response if more virulent strains than BCG were used, as increased secretion of factors has been observed with infection by more virulent mycobacterial strains.(Engele et al. 2002; Polena et al. 2016) Thus, these results support the use of our microscale granuloma model for studies of soluble factor signaling involving the granuloma layer, and its broader use for mycobacterial infection cytokine studies.(Domingo-Gonzalez et al. 2016)

**Figure 3:**
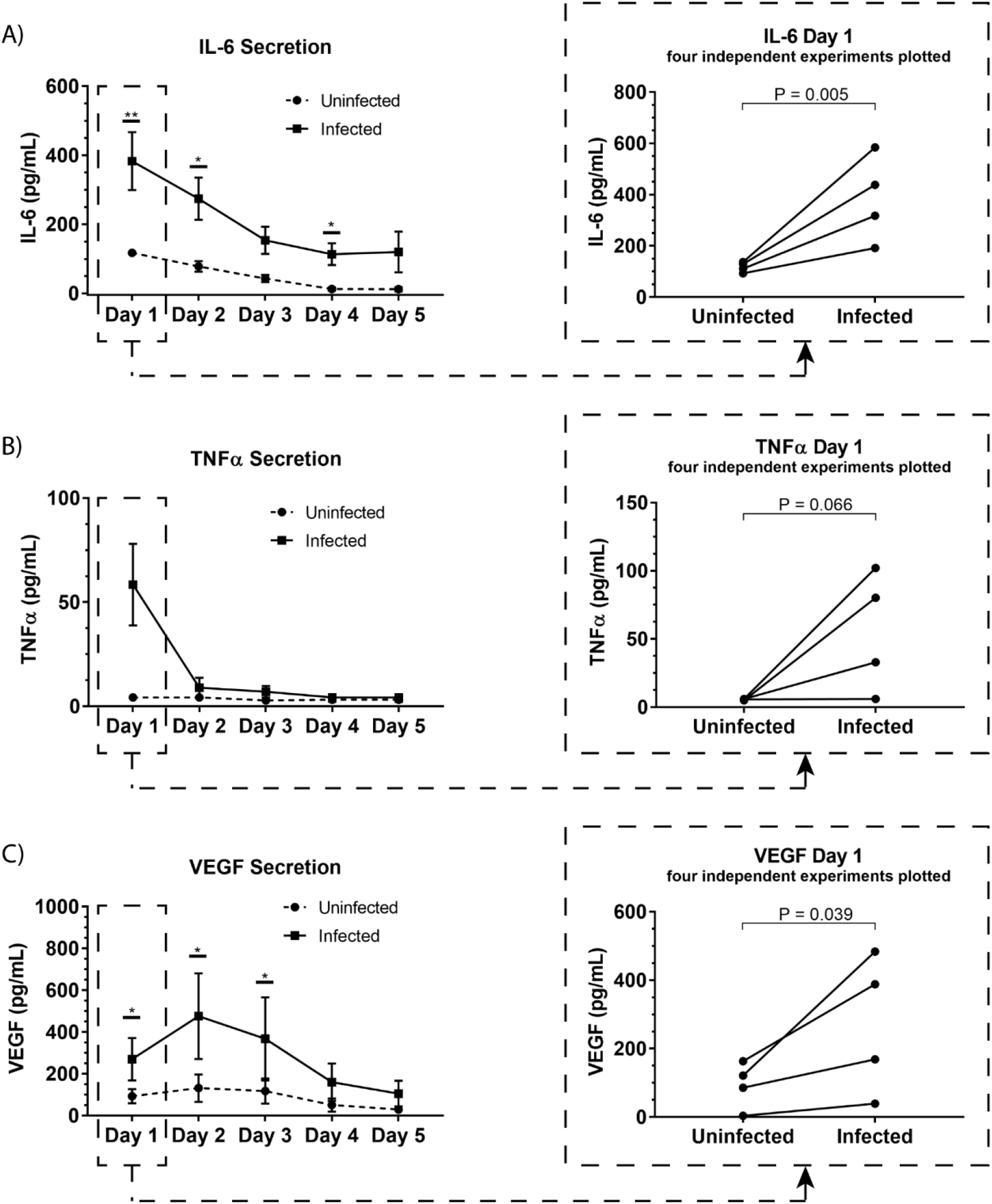
Soluble factor analysis of granuloma layer supernatants illustrates proinflammatory profile of infection model. Infection with BCG causes significant increases in secretion of (A) IL-6 and (C) VEGF in the Stacks platform, and shows an increasing trend in (B) TNFα secretion Day 1 p.i. Secretion of IL-6, TNFα, and VEGF decreases over time in infected granuloma layers following infection, corresponding with the formation of aggregates starting around Day 3 p.i. For Day 1-5 p.i., media was replaced daily and supernatant collected from each day was analyzed. Each point represents pooled supernatant samples from 24 technical replicates from n=4 independent experiments (each of the four independent experiments is plotted as a separate point in the plots on the right for the Day 1 data). Error bars on left plot: SEM. *P < 0.05; **P < 0.01; Ratio paired *t*-test.

### BCG Granulomas Modulate Endothelial Structure Morphology In Vitro

As a proof of concept to demonstrate the use of our microscale granuloma model in the Stacks platform, we developed a coculture system containing an established angiogenesis model(Theberge et al. 2015b; Yu et al. 2019; Lonza 2018; Koh et al. 2008; Sarkanen et al. 2011) that can be used to probe granuloma-associated angiogenesis.(Osherov and Ben-Ami 2016; Polena et al. 2016; Oehlers et al. 2015; Datta et al. 2015) Mycobacterium-mediated angiogenic processes and granuloma vascularization result from the secretion of proangiogenic factors by the infected immune cells that compose the granuloma and play a complex and evolving role during the course of infection.(Torraca et al. 2017; Polena et al. 2016; Oehlers et al. 2015; 2017) Extensive work has been conducted on understanding the role of angiogenesis in granuloma outcome, finding that inhibition of VEGF and other signaling pathways reduces pathogenicity and dissemination of infectious mycobacteria(Polena et al. 2016; Oehlers et al. 2015; Harding et al. 2019) while normalizing surrounding vasculature, improving small molecule delivery, and decreasing hypoxia within the granuloma.(Datta et al. 2015) Similarly, we observed increased secretion of VEGF from infected cells within our microscale granuloma model (Figure 3C), and therefore sought to illustrate one potential use of our platform as a complimentary tool for studies examining this infection-mediated process.

To establish this granuloma-vasculature model coculture system, we created a second Stacks layer containing a floor, enabling culture of human endothelial cells on a hydrogel plug (Matrigel) while retaining the ability to be placed in soluble factor signaling contact with the granuloma layer. We selected an *in vitro* model of angiogenesis that has been extensively used to screen angiogenic stimulants and inhibitors, wherein human endothelial cells cultured in well plates self-assemble into tubule-like networks and demonstrate cell sprouting and branching, and adapted it for our endothelial layer.(Bishop et al. 1999; Lonza 2018; Koh et al. 2008; Sarkanen et al. 2011) In order to examine the influence of the soluble factor profile from the granuloma layer on the endothelial layer, we independently established the granuloma layer and the endothelial layer under their optimized culture conditions; the modularity of the Stacks platform enables both layers to be cultured in their optimal conditions (and in separate incubators) without risk of cross contamination prior to joining the layers. The granuloma layers were infected and incubated 24 hours prior to clicking with the endothelial layer, and the endothelial layers were seeded 2 hours prior to allow the cells to self-assemble into a tubule-like network.(Lonza 2018; Sarkanen et al. 2011) Separate layers were then stacked together and connected by a bridge of shared media to allow passage of factors between the two layers. After 16 hours of signaling, we separated the layers and fixed, stained, and imaged the vasculature to analyze its morphology as a result of coculture with the granuloma layer (Figure 4). We found that after 16 hours of coculture, endothelial cells in contact with infected granuloma layers (+BCG) formed thinner, centralized structures with diffuse cell sprouts extending from the center, whereas endothelial cells in contact with uninfected control layers (-BCG) formed wider and larger structures that retained some interconnected networks (Figure 4). We quantified these morphological differences through measurement of their tubule index, a metric that measures the ratio of the endothelial structure perimeter to the endothelial structure area and can be used to discern between endothelial structure morphologies (e.g., tubule network, single tubule, islands/clusters)(Theberge et al. 2015a), and found significant morphological differences between endothelial layers cocultured with infected granuloma layers (+BCG) when compared uninfected granuloma layers (-BCG) (Figure 4). The morphological change is likely the result of the increased secretion of factors from BCG-infected MDMs, such as VEGF (Figure 3C); however, it is possible that additional factors we did not quantify are also contributing to the differences in endothelial morphology. Ultimately, the induction of morphology changes in the endothelial layer as a result of coculture with different model granuloma layers illustrates the ability of the granuloma layer to signal with a neighboring layer representing microenvironmental components.

**Figure 4:**
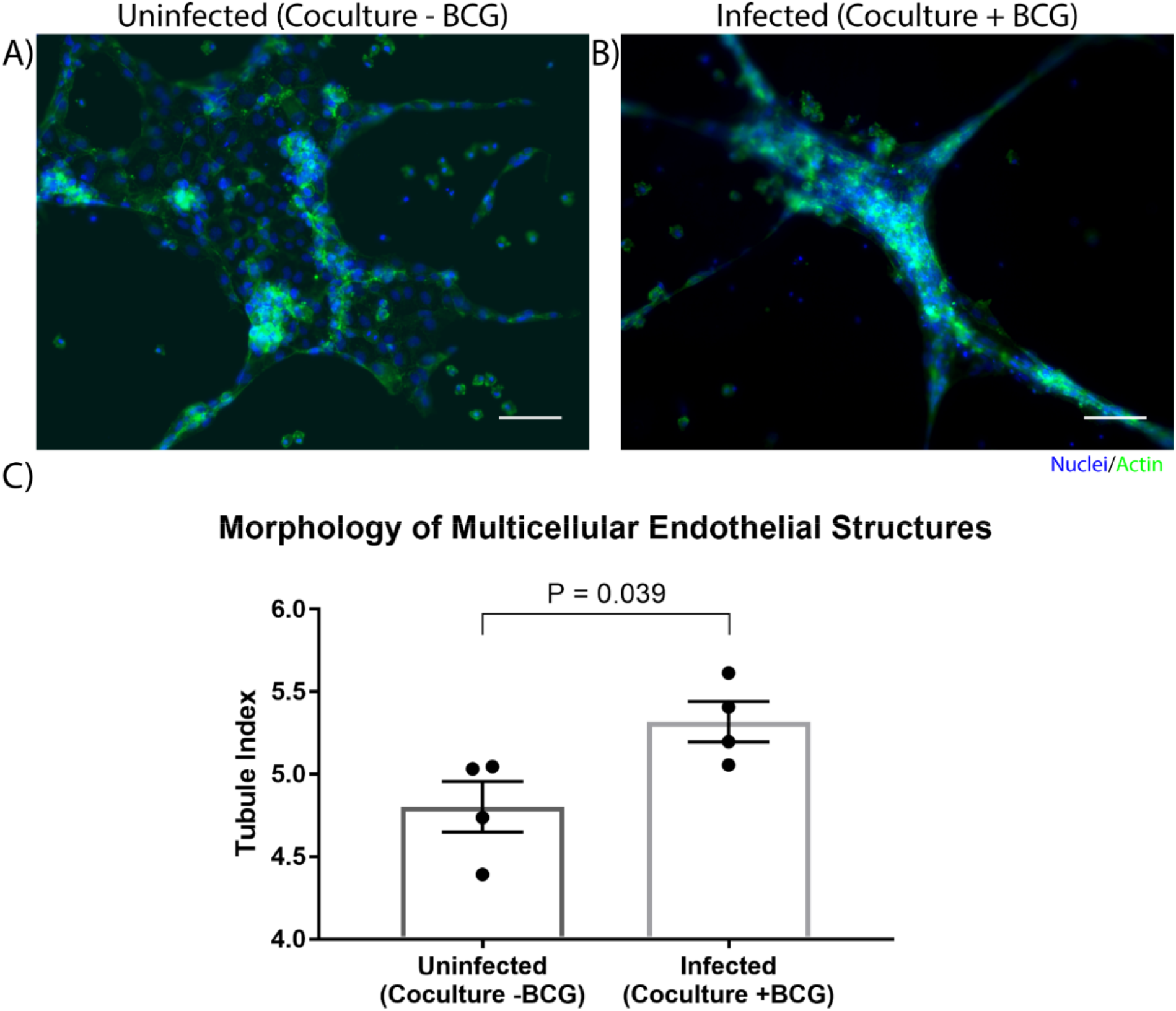
Coculture of endothelial layer with granuloma layer impacts morphology of multicellular endothelial structures. Representative images of an endothelial layer cocultured with an uninfected (-BCG) layer (A) or infected (+BCG) layer (B) for 16 hours, illustrating the difference in endothelial morphology as a result of coculture. C) The tubule index (perimeter/area ratio) was used to quantify the morphology of endothelial structures and shows a significant difference between coculture conditions, demonstrating the ability of the model granuloma layer to signal with the endothelial layer. Scale bar: 100 μm. Error bars: SEM. P < 0.05, unpaired *t*-test. Each point represents the average ratio of an independent experiment (n=4 independent experiments are plotted).

As we chose to connect the layers at Day 1 p.i. to correlate with increased levels of VEGF and cytokine secretion (Figure 3), we expect our model and results would mimic an earlier time point in infection, when the mycobacteria-infected cells are still secreting proinflammatory factors to recruit other immune cells and the vasculature. Depending on user queries, these layers can also be stacked at more advanced time points to study the effect of a later soluble factor profile on the surrounding vasculature, illustrating a key advantage of the modularity of the Stacks system and the flexibility in timing to combine the layers. Further, the ability of the model granuloma layer to communicate with the vasculature and modulate the endothelial morphology demonstrates the usage of this platform for signaling studies between our infected granuloma layers and microenvironment components. The ability to create two independent layers that are in soluble factor signaling contact enables users to isolate the effects of the soluble factors from the granuloma layer on its surrounding microenvironment without interference from juxtacrine signaling or physical interactions.

## Conclusion

Microenvironmental effects, such as soluble factor signaling, play a vital role in granuloma outcome, as the immune system attempts to contain and impede pathogenic mycobacterium from manipulating its environment in favor of its survival.(Ramakrishnan 2012) For example, induction of angiogenic processes by Mtb in the microenvironment surrounding a granuloma have been linked to pro-pathogen outcomes(Osherov and Ben-Ami 2016; Oehlers et al. 2015), while treatment with anti-angiogenic factors can be a potential treatment option to improve the effects of existing drug regiments.(Datta et al. 2015) To further understand the signaling environment and timeline of these phenomena, we created a novel *in vitro* granuloma model that can be used to study soluble factor signaling between the granuloma and its surrounding microenvironment. As a proof of concept model, we created a two-layered modular microfluidic system containing a mycobacterial infection and an endothelial vasculature model that can be used to compliment *in vivo* granuloma and angiogenesis models.(Polena et al. 2016; Oehlers et al. 2015; 2017; Datta et al. 2015) We demonstrate creation of a viable mycobacterial infection using human blood-derived immune cells and BCG within a 4 μL suspended collagen plug, and validate the secretion of proinflammatory cytokines associated with mycobacterial infection. Further, we provide a model coculture system that can be used to probe granuloma-associated angiogenic processes *in vitro.* The modularity and microscale size of our model enables users to add or remove additional cell types at various time points, and to utilize limited cell populations, such as patient cells or rare immune cells, and valuable reagents, such as antibodies or expensive drugs. Additionally, the design of the platform and fabrication method supports the creation of arrays to test various culture conditions/treatments and customizability to introduce additional functionality (e.g., flow), as well as easy integration with BSL3 laboratory workflows as the device is disposable and fits inside common cell culture materials (e.g., petri dish, OmniTray™, etc.). Further development of our system includes the addition of adaptive immune cells (e.g., T cells) and introduction of parenchymal and stromal cells (e.g., epithelial cells, fibroblasts) to model signaling interactions in the pulmonary environment, as well as the use of virulent *M. tuberculosis* to induce more *in* vivo-like responses. Ultimately, we created a tractable and customizable mycobacterial infection model that can be utilized by other researchers to examine the various signaling components of a complex tuberculosis infection.

## Materials and Methods

### Stacks Device Fabrication

Stacks devices(Yu et al. 2019) were fabricated from either polypropylene (PP) (granuloma layer) or polystyrene (PS)(endothelial layer)(Supplementary Figure 1, Table 1). PP devices were injection molded (ProtoLabs, Maple Plain, MN, USA) and were flattened using a bench top manual heated press (#4386, Carver Inc., Wabash, IN) for 1 h at 110°C (protocol in Supplementary Information 1). After flattening, devices were cleaned with isopropanol (IPA) sonication for 1 h to remove any fabrication residue or contaminants, and then rinsed twice with fresh IPA before drying with compressed air. Prior to use, devices were UV-sterilized for 30 min in a biosafety cabinet. PS devices containing a floor were designed using Fusion360 CAD software (Autodesk, San Rafael, CA, USA) and milled using a Datron Neo CNC mill (Datron Dynamics Inc., Livermore, CA, USA). PS devices and bottoms (floors, 53 mm x 53 mm) were milled from 1.2 mm thick PS sheets (Goodfellow Corp, Coraopolis, PA, USA). To attach the floor to the PS Stacks layer, floors were solvent bonded to the bottom of the Stacks layer using acetonitrile at 75°C for 10 min, followed by 15 min at 75°C to allow excess acetonitrile to evaporate. Solvent-bonded devices were then sonicated in IPA for 80 min and 70% ethanol for 15 min. Devices were then soaked in sterile deionized water for a minimum of 3 hours, dried with compressed air, and UV-sterilized for 30 min prior to use. Device holders were designed on Solidworks CAD software (Solidworks Corp., Waltham, MA, USA), converted into G-code using SprutCAM CAM software (SprutCAM, Naberezhnye Chelny, Russia), and milled using a Tormach PCNC Micromill (Tormach Inc., Waunakee, WI). Device holders were fabricated from 4 mm thick PS and were sonicated in IPA for 1 h and UV-sterilized for 30 min prior to use. All device and device holder design files are included in the SI (Supplementary Table 1).

To prevent evaporation of microliter volumes of samples, culture platforms were placed in a Nunc Omnitray™ (ThermoFisher) which was then placed in a bioassay dish (#240835, ThermoFisher) for secondary containment; both the Omnitray and bioassay dish were filled with sacrificial water droplets (1 mL and 5 mL, respectively) to create a humidified environment around the platform, and all cultures were then placed into a water-jacketed incubator.

### Cell Culture

Human peripheral blood mononuclear cells (PBMCs) were isolated from patient whole blood samples (Bloodworks NW, Seattle, WA, USA) using the Ficoll-Paque PLUS media density separation protocol (ThermoFisher, Waltham, MA, USA). Briefly, blood samples were diluted with PBS (Fisher Scientific) and layered atop the Ficoll-Paque. Tubes were then centrifuged for 20 min at 1900 RPM without brakes, and afterwards white blood cells at the plasma-Ficoll-Paque interface were collected and resuspended in PBS + 2% fetal bovine serum (FBS, ThermoFisher) + 1 mM ethylenediaminetetraacetic acid (EDTA, Gibco, ThermoFisher). Cells were then rinsed with subsequent centrifugation (10 min, 500g) and resuspension until the supernatant was clear. Isolated PBMCs were then cryopreserved in solution containing 90% heat-inactivated FBS (HI-FBS) (ThermoFisher) and 10% dimethyl sulfoxide (DMSO) (Sigma-Aldrich, St. Louis, MO) and stored in liquid nitrogen until use. To differentiate the PBMCs into monocyte-derived macrophages (MDMs), PBMCs were thawed and resuspended in serum free RPMI 1640 media (Gibco, ThermoFisher); following resuspension, PBMCs were seeded into a 6 well plate (#3516, Corning Inc., Corning, NY, USA) and allowed to incubate for 3 h at 37°C and 5% CO_2_ for monocyte adhesion. After 3 h, suspended cells were removed, adherent cells were washed once with 1X phosphate-buffered saline (PBS, Fisher Scientific), and then RPMI 1640 containing 10% HI-FBS, 2 mM L-glutamine, 25 mM HEPES, and 50 ng/mL macrophage colony-stimulating factor (M-CSF) (R&D Systems, Minneapolis, MN, USA) was added to each well. Media was changed on Day 3 and Day 6 post-seeding, and MDMs were used after Day 6.

mCherry-expressing *M. bovis* Bacillus Calmette-Guérin (BCG) (graciously provided by the Urdahl Lab, Seattle Children’s Research Institute, WA, USA) was cultured in Middlebrook 7H9 broth (ThermoFisher) containing 10% Middlebrook ADC enrichment supplement (ThermoFisher), 0.003% Tween-80 (Fisher Scientific), and 50 μg/mL hygromycin B (Sigma-Aldrich). A lower Tween-80 concentration had to be used to ensure that the surfactant did not interfere with the microfluidic media pinning in the Stacks device (Supplementary Information 2). BCG was cultured at 37°C and 170 RPM to an OD of 0.7-1.0 for use, passed through a 27G needle to break up aggregates, and diluted to a working concentration for the experiment. BCG was used for all experiments as it can be used within a BSL2 facility, and we did not have access to virulent mycobacterium strains nor a BSL3 facility to perform these experiments.

Human umbilical vein endothelial cells (HUVECs) (Lonza, Basel, Switzerland) were cultured in completed EGM-2 media and maintained at 37°C and 5% CO_2_ until 80-90% confluence. Passage 5-7 cells were used.

### 3D Granuloma Assay

MDMs were differentiated for a minimum of 6 days prior to use, and BCG was grown to an OD of 0.7-1.0 for use. MDMs were rinsed once with 1X PBS and detached with enzyme-free Cell Dissociation Buffer (#13151014, Life Technologies, ThermoFisher) at 37°C for 5 min and vigorous pipetting. Detached cells were neutralized with complete RPMI 1640 media, counted, and resuspended at a density of 4×10^7^ cell/mL. BCG was vortexed for 30 seconds, vigorously mixed via pipetting, and then diluted into 4 mL of complete RPMI 1640 media for a final concentration of 2×10^6^ BCG/mL. An extracellular matrix (ECM) mix containing 80 μL 3 mg/mL type I bovine collagen (Advanced Biomatrix Inc., Carlsbad, CA, USA), 10 μL 10X HEPES buffer, 7 μL deionized H_2_O, 3 μL 0.5N NaOH, and 2.25 μL 1 mg/mL human fibronectin (Sigma-Aldrich) was mixed and stored on ice until use. To prepare the suspended cell-laden collagen plugs, 25 μL of MDMs (at 4×10^7^ cells/mL) and 25 μL of BCG (at 2×10^6^ BCG/mL) or 25 μL of complete RPMI 1640 was added to the ECM mix for a final volume of 152.25 μL and an MOI of 0.05. After mixing, 4 μL of the cell-laden ECM mix was added to each well in a PP device and allowed to gel at 37°C and 5% CO_2_ for 2 h. After gelation, 8 μL of RPMI 1640 containing 15% HI-FBS and 25 mM HEPES was added atop each well and returned to the incubator. Media was changed daily for the entirety of the experiments. For all monoculture infection experiments (Figures 2 and 3), MDMs were stained with CellTracker Green CMFDA dye (ThermoFisher) according to manufacturer’s protocols. Briefly, MDMs were rinsed with 1X PBS, and 10 μM CellTracker Green in serum-free media was added for 30 min at 37°C, washed once with 1X PBS, and then incubated with complete media for 10 min prior to further processing or use.

### 2D Angiogenesis Assay

Prior to seeding, PS Stacks devices containing a floor were chilled for 5 min at −80°C and then kept on ice for Matrigel seeding. 3μL of Matrigel (8.6 mg/mL) was added to each well and devices rested on ice 30 min. The Matrigel was then polymerized for 30 min at 37°C. After polymerization, 3 μL of HUVECs were added atop the Matrigel for a final concentration of 1,650 cells per well (5.5×10^5^ cells/mL). Cells were allowed to adhere and self-assemble for 2 h at 37°C and 5% CO_2_. Media was then aspirated and replaced with 2 μL of EGM-2 + 10% FBS coculture media.

To stack the layers, media was first removed from the model granuloma layers. Model granuloma layers were then placed atop the endothelial layers (containing 3 μL of media) and 7 μL of coculture media was placed atop each well of the stacked granuloma layers, to allow for feeding of both layers. Layers were separated by a thin layer of tape along the sides of the devices to ensure reproducible separation. Stacked devices were then placed in the incubator at 37°C and 5% CO_2_. After 16 h, devices were inverted and separated, fixed with 4% paraformaldehyde for 30 min at 25 °C, and stained as specified under Imaging.

### Imaging

To validate granuloma layer formation within our platform, MDMs were prestained with CellTracker Green CMFDA dye prior to infection and seeding into the Stacks. For imaging, the Stacks platform was placed on a 50 mm x 75 mm glass coverslip (#260462, Ted Pella Inc., Redding, CA, USA) and placed into an OmniTray™ with a 45 mm x 70 mm rectangle cut out of the bottom. Fluorescent images were obtained on a Zeiss Axiovert 200 coupled with an Axiocam 503 mono camera (Carl Zeiss AG, Oberkochen, Germany). For confocal imaging, all wells were fixed with 4% paraformaldehyde for 1 h at 25°C and then covered with PBS; the same imaging protocol was then followed as above. Fluorescent confocal images were obtained on a Leica TCS SP5 II Laser Scanning Confocal Microscope (Leica Camera AG, Wetzlar, Germany), with a z-depth of 66.47 μm and step size of 1.01 μm. Images obtained with the Zeiss Axiovert 200 were analyzed with Fiji (ImageJ), and images obtained with the Leica TCS SP5 II were analyzed with Leica LAS X software (Leica).

For vasculature layer imaging, cells were permeabilized with 0.5% Triton-X 100 for 30 min and then stained with Phalloidin 488 (1:50) (Molecular Probe A12379) and Hoechst nuclear stain (1:1000) (Molecular Probe H1399) overnight. After overnight incubation, cells were rinsed thrice with 0.2% Triton-X 100 for 10 min each and then covered with PBS for imaging. Devices were placed on a 75 mm x 50 mm glass coverslip (#260462, Ted Pella Inc., Redding, CA, USA) and placed in an Omnitray™ with a 45 mm x 70 mm rectangle cut out of the bottom. Fluorescent images were obtained on a Zeiss Axiovert 200 coupled with an Axiocam 503 mono camera (Carl Zeiss AG, Oberkochen, Germany). Images obtained were analyzed with Fiji (ImageJ).

### Angiogenesis Morphological Analysis

To analyze differences in morphology in the endothelial layer we used default functions of Image J (Fiji) to quantify the tubule index (perimeter/area ratio)(Theberge et al. 2015a). The image analysis procedure is described in the methods and SI of Theberge et al.(Theberge et al. 2015a), a summary is included here. For quantification, the 8-bit image containing the phalloidin channel (488) was selected and a threshold (Huang Dark) was applied to get the outline of the endothelial culture and saved (Image 1). The “Fill Holes” function was applied and saved (Image 2), and then Image 2 was subtracted from Image 1 to generate an image containing the holes/gaps in the endothelial morphology (Image 3). The total area of the phalloidin stain was then calculated by subtracting the area of Image 3 (holes) from Image 2 (fill holes), and the perimeter of the stain was calculated by adding the perimeter of Image 2 and Image 3. The area and perimeter were measured using the function “Analyze Particles”, with the following parameters: size: 25-infinity (include pixel units), circularity: 0.00-1.00. The ImageJ macro for this process is included in the supplementary materials and is based off the macro used in Yu et al.(Yu et al. 2019)

Exclusion criteria for experimental artifacts that interfered with the imaging technique was developed to exclude wells unable to be analyzed with this technique. For example, if a collagen plug from the model granuloma layer detached and fell into a endothelial layer well during separation of the layers, the endothelial morphology was obscured by signal from the MDMs in the collagen plug and therefore unable to be accurately measured. All images were assigned a randomized code and individuals not involved in the project determined which images to exclude base on established criteria. Images were then analyzed according to the measurement protocol above.

### Cytokine Analysis

Cytokine analyses were performed with a custom Luminex ProcartaPlex multiplex assay for IL-6, TNFα, and VEGF (ThermoFisher). Supernatant samples were collected and pooled from a single device (n=24 wells, 8 μL/well) during daily media changes and stored at −80°C until use. Samples were collected from the same device over a five-day period, such that each sample contains cytokines secreted within 24 h of collection (i.e., Day 2 p.i. cytokine levels indicate what was secreted between Day 1 and Day 2 p.i., etc.). For analysis, samples were thawed on ice and analyzed according to the manufacturer’s protocols for Luminex multiplex assays. Samples were analyzed on a Luminex 100/200 System instrument with xPONENT software (Fred Hutchinson Cancer Center Core Facility). All results were analyzed and visualized using Prism 7 software (GraphPad Software, San Diego, CA, USA).

## Supporting information

Berry et al. SI Final

ImageJ Macro

Design Files

## Acknowledgements

We would like to thank Dr. Shahin Shafiani (Urdahl Lab) for providing the mCherry *M. bovis* BCG strain, Dr. Jason Yu for providing help on the Stacks platform, Tammi van Neel and Tianzi Zhang for image review, Ashley Dostie for obtaining the Stacks image, and Erik Layton (Seshadri Lab) for PBMC isolation training. We acknowledge the support of the Biochemical Diagnostics Foundry for Translational Research supported by the MJ Murdock Trust. This work was supported by NIH R35GM128648, the University of Washington, the Mary Gates Endowment, and by the National Science Foundation Graduate Research Fellowship Program under Grant No. DGE-1256082 (SBB). Any opinions, findings, and conclusions or recommendations expressed in this material are those of the author(s) and do not necessarily reflect the views of the National Science Foundation.

## Author Contributions

SBB and ABT conceived of the project. SBB and MSG performed all experiments and data analysis. SBB and MSG wrote the manuscript, and XS, CS, and ABT advised the project and edited, revised, and provided feedback on the manuscript. All authors reviewed and approved the final manuscript.

