## Supplementary material for "A modular microscale granuloma model for immune-microenvironment signaling studies *in vitro*": Berry et al. SI Final

The supporting information for “A modular microscale granuloma model for immune-microenvironment signaling studies *in vitro*” includes information that readers might find useful for adapting this platform for their own research and laboratory setups. We include detailed technical schematics of our Stacks platform, the code for the ImageJ macro we adapted for analysis of endothelial morphology, device preparation protocols, experimental optimization considerations, and design files for the devices used in this manuscript.

A. 3D Injection Molded Device for Model Granuloma Layer

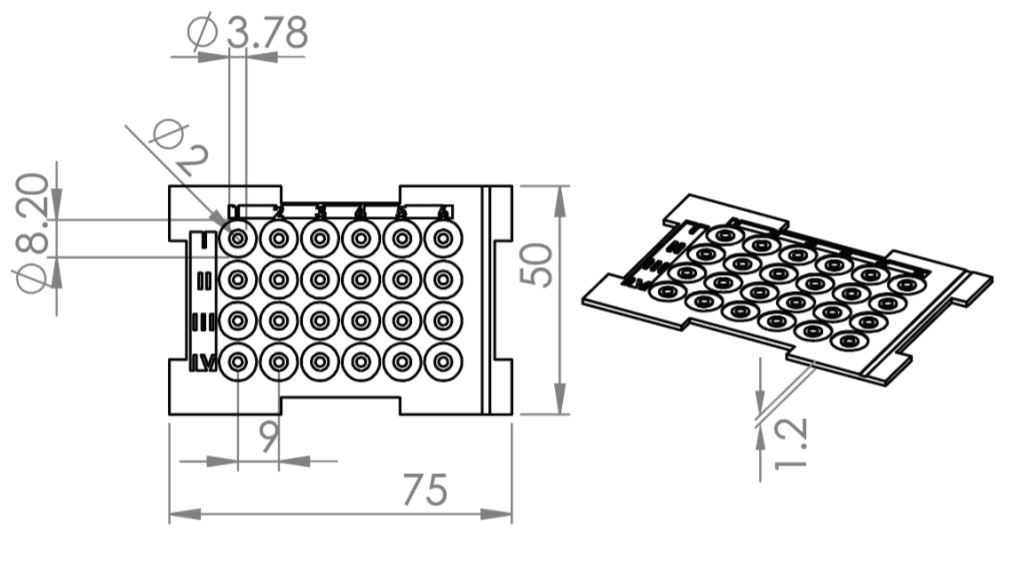

B. CNC-Milled Device for Endothelial Layer

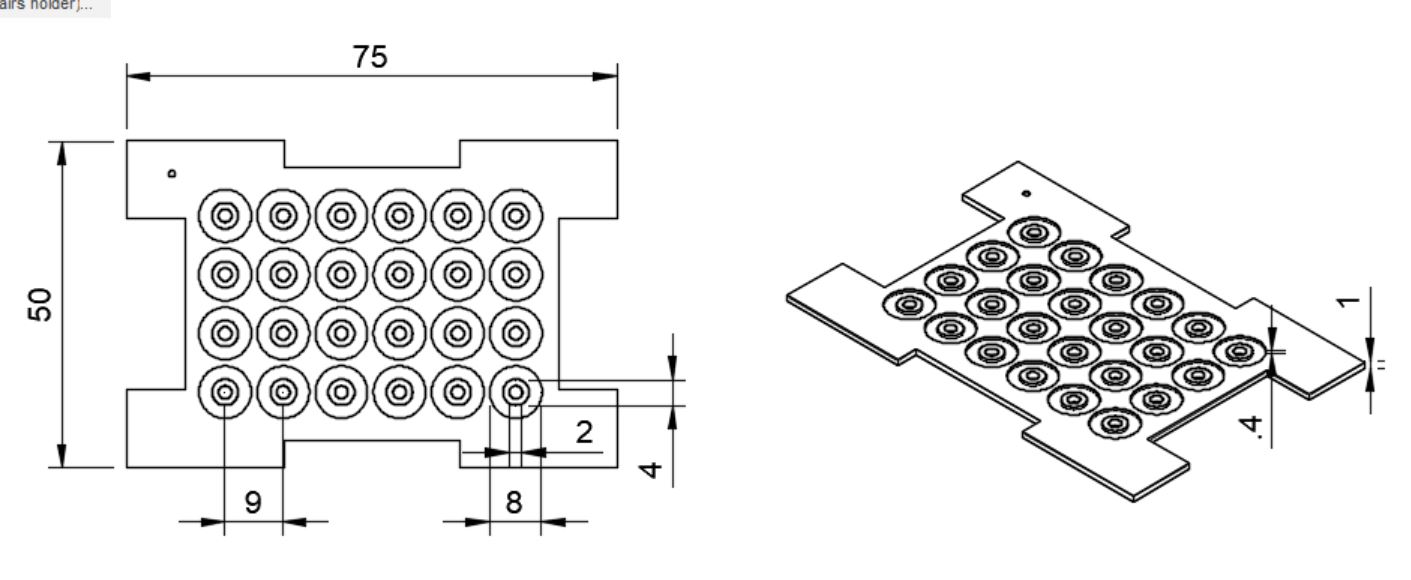

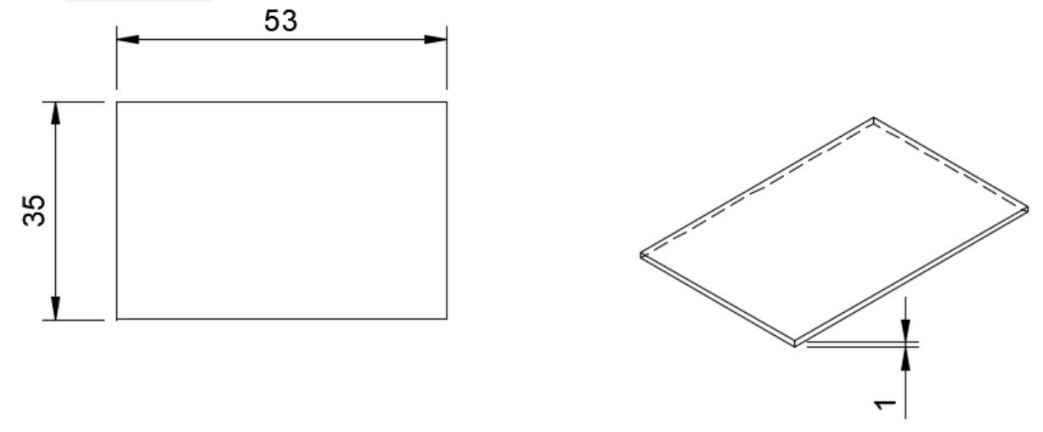

**Supplemental Figure 1:** Detailed Device Schematics and Dimensions. A) The injection molded layer used in Figures 1-4 for the model granuloma. Dimensions and figure reproduced from Yu et al.^1^, Supplemental Figure 3. B) The endothelial layers containing a floor used in Figure 4 was milled using CNC bonding with a solvent-bonded floor. Dimensions labeled in millimeters.

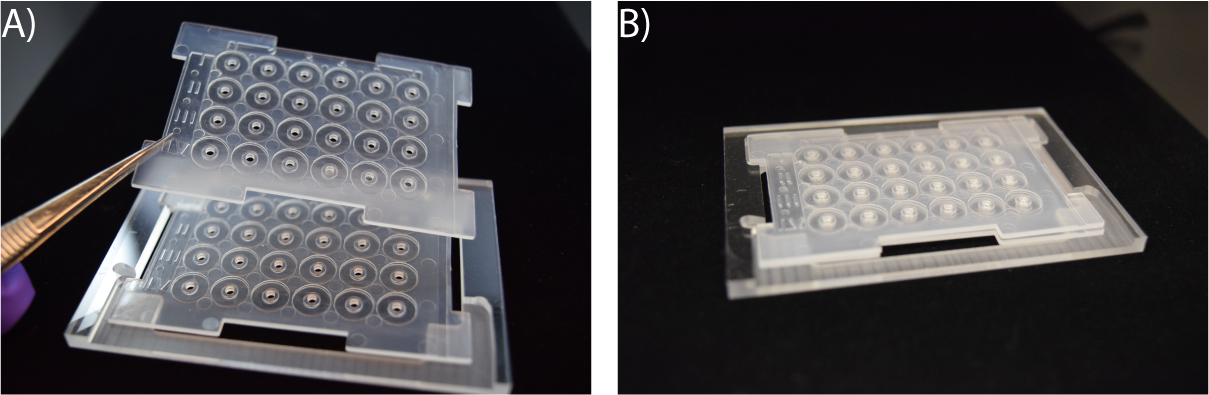

**Supplemental Figure 2:** Stacks devices placed inside of CNC-milled device holders. A-B) Layers can be placed directly into the holders to prevent contact between the open suspended culture wells and the floor, as well as to keep the devices in place during culture and aligned once vertically stacked.

**Supplementary Information 1: Polypropylene (PP) Device Flattening Protocol**

Protocol is for use with a Carver Bench Top Standard Heated Press (Model #4386). This protocol can be adapted for use with alternative heated presses with model-specific changes.

1. Preheat platens to 110°C (this takes ≈ 20 min to heat up and stabilize, as the temperature fluctuates).
2. Prepare a stack of devices (4-5 devices) and carefully align the stack to avoid deformation.
3. Place Kapton® polyimide film (#2271K2, McMaster Carr) on the top and bottom platens and place the stack of devices between the films (to avoid direct contact between the devices and the platens).
4. Turn on the pressure sensor.
5. Pump the hot press handle until contact is just made and the devices are flattened.

- The devices stacks should not be compressed at all (i.e., the pressure sensor should read "0 psi" or fluctuate between “0-10” psi), but the devices should be in contact with the platens and flattened.

6. Let the stacks flatten for 60 min at the above temperature and pressure.
7. After 60 min, turn off the platen temperature and let the temperature drop down to ≈30°C.

- Do not use coolant to lower the temperature as this causes the temperature to drop too rapidly and can cause deformation. This cooling process takes 1.5-2 h.

8. Once the temperature has dropped below 30°C, turn on the coolant circulator and run the coolant for 5 min to completely cool the platens.
9. Turn off the coolant, release the pressure, and remove the devices from the platens.

**Supplementary Information 2: Culture Optimization for Miniaturization of Mycobacterial Infection**

| **Condition** | **Conventional Macroscale Protocol** | **Optimized Microscale Protocol** | **Reason for Optimization** |
| --- | --- | --- | --- |
| Fetal Bovine Serum Concentration^2^ | 10% | 15% | Adaptation for microculture leads to decreased volume of media per cell; increased [FBS] provides greater amount of serum to culture. |
| Media Buffer Concentration^2^ | Variable (0-25 mM HEPES) | 25 mM HEPES | Adaptation for microculture leads to decreased volume of media per cell; increased [HEPES] provides greater buffering capacity of media to counter waste accumulation. |
| Culture Feeding Frequency^2^ | Every 2^nd^ or 3^rd^ day | Daily | Adaptation for microculture leads to decreased volume of media per cell; increased media changing frequency prevents nutrient depletion and waste buildup. |
| Multiplicity of Infection (MOI) | 0.1^3,4^ | 0.05 | Adaptation to microscale decreases the total volume of space in the culture well and increases probability of BCG-recognition by monocyte-derived macrophages; decreased MOI results in reproducible and consistent aggregate formation in each well with sufficient uninfected MDMs remaining to aggregate around the infected cells. |
| Tween-80 Concentration | 0.05% | 0.003% | Adaptation for the Stacks platform requires a Tween-80 concentration below the critical micelle concentration to maintain capillary pinning and functionality of Stacks devices (see below). |
| Device Material | Polystyrene (PS) | Polypropylene (PP) | Adaptation for use with immune cell and mycobacterial media requires an increase in the contact angle to maintain capillary pinning of media for cultures. |

Optimization of the Tween-80 concentration for culture of *M. bovis* BCG was required for compatibility with the Stacks platform. Traditionally, mycobacteria are cultured in varying concentrations of surfactant (commonly Tween-80) to prevent clumping of mycobacteria during culture. However, the presence of surfactant within the Stacks platform interferes with the capillary pinning necessary for maintenance of cell cultures and stacking of multiple layers. This is due to the decrease in interfacial tension between the culture media and the device surface caused by the presence of surfactant, effectively decreasing the contact angle of the media on the surface. In order to alleviate the effects of surfactant on the capillary pinning, the concentration of surfactant must be well below the critical micelle concentration (CMC), which is 0.0013% w/v^5^, in the final media. Therefore, we cultured the BCG at a concentration of 0.003% w/v Tween-80, which yields a final concentration (after all dilutions and mixing) of 0.0000025% w/v Tween-80 in the collagen plug; this concentration was selected as it was the highest concentration of Tween-80 we could use without loss of capillary pinning on the surface. However, to decrease the aggregation of BCG within our model system, the BCG were vortexed, vigorously pipetted, passed through a 27G needle to disperse aggregates, and then allowed to settle for ≈ 1 min before aliquoting from the top of the culture for use.

**Supplementary Information 3: Culture Optimization of Endothelial Layer**

To miniaturize the *in vitro* angiogenesis model^6-8^ we adapted for this layer and to ensure functionality, we optimized certain components of the culture system within the Stacks platform. The first condition optimized was the seeding density; too high of a seeding density results in formation of a confluent monolayer or islands of cells on the surface of the Matrigel, whereas too low of a seeding density results in a dispersed culture that does not form endothelial connections. Therefore, the seeding density was calculated from previously established protocols using 48 well plates^7^ to obtain a similar ratio of cells to area in the Stacks well (1,650 cells/well of the Stacks layer). We observed that concentrations greater or less than 1,650 cells/well resulted in inconsistent tubule or network formation.

Additionally, the volume of Matrigel within each well was optimized. The final volume of 3 μL was selected to allow complete and reproducible coating of the bottom of each well and to limit the effect of the meniscus on cell localization (i.e., cells aggregating towards the middle of the well/bottom of the meniscus) and cell visualization. Lower volumes of Matrigel (< 3 μL) resulted in uneven coating of the surface of the well floor, causing a non-uniform surface wherein cells adhered to both Matrigel and exposed polystyrene on the floor of the well, leading to altered morphology. Higher volumes of Matrigel (> 3 μL) caused an exaggerated meniscus effect, resulting in cell aggregation in the center of the well. In cases where the volume of the well was completely filled with Matrigel (> 3.6 μL), we observed cell growth both on the Matrigel and on the polystyrene pinnng ridge surrounding the well, resulting in varying surface-dependent morphologies (i.e., cells displayed different morphologies on the polystyrene and the Matrigel).

**Supplementary Table 1: Files Included in Supplemental Information**

| **Figure** | **File Name** |
| --- | --- |
| Figure 1-4 – Injection Molded Device^1^ (3D) Model Granuloma Layer | *3D Injection Molded Layer.f3d* |
| Figure 1-4 – CNC-Milled Device Holder | *Stacks Device Holder.f3d* |
| Figure 4 – CNC-Milled Device (2D) Endothelial Layer | *2D CNC Milled Layer.f3d*  *2D CNC Milled Floor.f3d* |
| Figure 4 – ImageJ Macro for Analysis | *ImageJ Macro Image Preparation and Analysis.txt* |

The file for the injection molded device was reproduced from Yu et al.^1^
